# Multifunctional phosphate based nanoparticles as a platform for imaging, targeting and doxorubicin delivery to human breast cancer CD44^+^ cells

**DOI:** 10.1101/2021.11.08.467718

**Authors:** Priscila Izabel Santos De Tótaro, Betânia Mara Alvarenga, Diego Carlos dos Reis, Thaís Maria da Mata Martins, Anderson Kennedy Santos, Rodrigo Ribeiro Resende, Geovanni Dantas Cassali, Alfredo Miranda Góes, José Dias Corrêa Júnior

## Abstract

Functionalized nanostructured systems can be used for imaging and drug delivery for anti-tumor therapy, including breast tumors. This is a more efficient approach that offers reduced systemic side effects compared to conventional diagnostic and chemotherapy methods. Multifunctional nanoparticles are potential tools in the diagnosis, location tracing and kill tumor cells through a less invasive manner. Functionalized phosphate-based nanoparticles are capable of encapsulating, or may be associated, with fluorescent probes. In this study, we synthesize a nanoparticle phosphate-based composite (NPC) and functionalize it with poly-ethylene glycol (PEG), hyaluronic acid (HA), the fluorescent probe rhodamin 6G (R6G) and the antimitotic doxorubicin (DOX). We focused on targeting human breast cancer cells reporting the biological effects of functionalized NPC on them. NPC and NPC formulations containing PEG, HA, and R6G did not cause cell viability reduction on MCF-7 and MDA-MB-231 cell lines. The cellular internalization of NPC was quantified by real-time in vitro observation, and confirmed by electron microscopy techniques. Intracellular NPC distribution is detected in the cytoplasm and nucleus of tumor cells by confocal fluorescent images. The percent association of doxorubicin to NPC matrix was approximately 18% and NPC formulations associated with doxorubicin led to a significant reduction in cell viability in MDA-MB-231 and MCF-7 cells. This data suggest the potential use of NPC as a non-cytotoxic platform for association with functional ligands to selective targeting breast cancer cells. NPC use can be also explored in drug delivery to cancer cells.

## Introduction

Cancer is a disease that due to its high mortality rate, is one of the major concerns in global public health, and accounted for almost 1,685,210 new cases in 2016 in USA. Although there are modern therapies available, little success has been realized when the disease has reached an advanced stage; additionally, some peculiar types of cancers have a very low cure rate. As such, cancer is still a major challenge for many medical researchers.^1^

Nanotechnology applied to cancer studies involves the use of drug delivery platforms and the targeting of tumour cells in vivo and in vitro.^2, 3, 4, 5^ Nanoparticles have unique and differentiated proprieties: are expected to accumulate in tumours due to enhanced permeation and retention (EPR) effect^6^ and possess a tunable surface that allows the development of passive or active targeting systems.^7^

Meanwhile, for being used as an anticancer or tumour detection agent, a nanoparticle could possess a stealth layer and a ligand for cancer reception. Poly-Ethylene-Glycol (PEG) is one of the most common ligands used for stealth function,^6^ due to its hydrophobicity and flexibility, ^8,9^ and for in vivo approaches, PEG enhances the nanoparticles’ lifetime circulation by preventing their interaction with plasma proteins and mononuclear fagocitary system cells.^10, 11, 12, 13^ Additionally, the targeting ligands must possess sufficient specificity in order for the receptors to be overexpressed on cancerous cells being minimally or normally expressed on healthy cells. These molecules must also have a high affinity to their receptors and possess the ability to induce receptor-mediated endocytosis.^6^ Active targeting systems to enhance the internalization efficacy of nano carriers in cancer management are very diverse and may involve a few different kinds of biological molecules.^1^ A strategy often used to target cancer cells is the nanoparticle functionalization using hyaluronic acid (HA),^14, 15, 16, 17^ a water-soluble and biodegradable polymer, fully distributed in the human body,^18^ and a natural ligand for CD44. CD44 is a trans membrane adhesion glycoprotein receptor, participating in cell–cell and cell–extracellular matrix interactions^19^ that is expressed in many cell lines like leukocytes and fibroblasts, participating in cell migration and lymphocyte homing,^20^ also being considered one of the most widely accepted cell surface markers for a variety of cancer cells, including human breast cancer cell lines MDA-MB-231 and MCF-7, in which is overexpressed.^21^ Therefore, the interaction of CD44 and HA can be used as a potential target for cancer diagnosis and therapy.^22, 23, 24,25^

Another great challenge in cancer research is the early detection of disease, which motivates the use of contrast agents, such as imaging markers that can be made specific to cell types,^26^ like breast cancer cells.^27,28^ Platforms for monitoring or imaging during diagnosis and therapy must involve a fluoroprobe, which retains its state of dispersion in physiological solutions or environments, has prolonged signal intensity, is able to accumulate in the regions of interest, and resorbs into the body after use. In this scenario, nanotechnology has brought to bioimaging, the capacity to improve fluoroprobe molecule efficiency in vivo.^29, 30^

By concentrating and protecting a marker for degradation, the association of probes with nanoparticles may turn analyses into being more sensitive.^8^ Thus, nanomaterials have been explored as synthetic scaffolds for imaging probes used in the detection and targeting tumours in cancer therapy.^7^

Preliminary tests conducted in our laboratory have demonstrated our ability to synthesize low-cost nanostructured phosphate composites (NPC). They were previously successfully tested as transfection agents in difficult-to-transfect-cells, including neonatal rat cardiomyocytes, rat dorsal root ganglion cells (DRG cells), human neuroblastoma cell line (U37311 cells), rat glial tumour cell line (C612 cells), ^31^ and fish spermatogonial stem cells.^32^ Moreover, they presented promising results for antimony (Sb) delivery on leishmania infected murine macrophage, thereby targeting and reducing chemotherapy in in-vitro assays.^33^

Thus, because calcium phosphate-based nanoparticles have previously reported good safety of healthy cells, low price, facility to syntheses, biocompatibility, biodegradability^34^ in this work, we aimed to use a nanoparticle phosphate-based composite (NPC) and functionalize it using PEG and HA (NPCF), rhodamin 6G a cytotoxic red organic dye (NPCFF) and doxorubicin (NPC-DOX) as an anti-mitotic complex. In this paper, we aim to describe the physicochemical proprieties and some biological roles of different NPC formulations and expect to reach to a cheap, nontoxic multifunctional and fluorescent platform in order to target and treat CD44^+^ human breast cancer cells. The data presented here provide evidences that NPC could be used as a platform for new products and therapeutic protocols with potential advantages over the methods currently applied.

## Experimental

### Synthesis of Nanostructured Phosphate Composites (NPC)

Nanostructured phosphate composites (NPC) were synthesized in the Laboratory of Chemical–Biological Interactions and Animal Reproduction, Morphology Department, ICB-UFMG, using a liquid medium and controlled pH solutions containing phosphate salts and metals of known biological activity. These compounds have been associated with crystalline state modifiers (Na_4_P_2_O_7_ - Cromoline; CaCl_2_-2H_2_O - Cromoline; MgCl_2_-6H_2_0-Synth). Using a system of semipermeable membranes (Spectrum Medical Industries, Inc.), NPCs were obtained. The experimental conditions and the preparation methodology of NPCs are in form of patent deposits registered under the numbers 102012032493-8 BR, and BR 102013032731-0. Post synthesis NPCs were isolated by centrifugation at 3500 rpm for 10 minutes, discarding the supernatant. The resulting precipitate was washed thrice with absolute ethanol (Merck), and following further centrifugation, it was oven dried at 60°C for 48 hours. Amounts of 1 mg of each NPC were placed in conical polypropylene microtubes and sterilized by gamma irradiation at a dose of 25 kGy, CDTN-UFMG.

### Functionalization of NPCs with Hyaluronic Acid (HA) and Polyethylene Glycol (PEG) (formation of NPCF)

A stock solution containing NPC (1 mg) was eluted in milli-Q water (1 ml) and homogenized with through a vortex for 5 minutes. To the resulting solution, were added hyaluronic acid (HA) (30 μL at 9.71 mg ml^-1^) (Galena Chemical and Pharmaceutical Ldta) and, after further homogenization, polyethylene glycol (PEG 400 – synth) (3 μL) at room temperature. All analyses were performed after a minimum of 15 minutes and a maximum of 24 hours following the end of functionalization.

### Functionalization of NPCs with rhodamin 6G (NPCFF)

Stock solutions of rhodamin 6G (R6G – Sigma–Aldrich) (1.0 mM) were eluted in water to carry out the adsorption test to be performed with NPC. NPC stock solution (1 mg ml^-1^ in milli-Q water) received 20 μL of R6G (1.0 mM). To the resulting system, 25 μL of hyaluronic acid (9.71 mg ml^-1^) and 3 μL of PEG 400 (NPCFF) were added sequentially.

### Characterization of Nanostructured Phosphate Composite (NPCs) and Functionalized Nanostructured Phosphate Composite (NPCFs)

#### Zeta potential and conductivity

To obtain the zeta potential of NPCs and NPCFs, Zetasizer Nano Series equipment (Malvern) were used. NPC stock solution (200 μL) and NPCF (1 mg mL^-1^ in milli-Q water) were added to 2 ml of distilled water. Readings for the resulting solution were taken and recorded after homogenization and stabilization of the sample.

#### Diameter distribution, morphology, and elemental composition through X-ray microanalysis

Aliquots of 10 μL NPC stock solution (1 mg mL^-1^ in milli-Q water) were deposited on a carbon coverslip (Termanox®), air-dried, vacuum metallized (10^−5^ mbar) with carbon in an evaporator (Hitachi Model HUS4G), and analyzed in scanning microscopes (FEI Quanta FEG 200 and JEOL JSEM6340) at the electron Microscopy Center of UFMG. The resultant images were acquired by secondary electrons, using magnifications between 5,000 and 50,000×. NPC diameters were obtained through the morphometric technique using Image Pro Plus 4.0 software. Diameter distributions were generated in GraphPad Prism 5.0 software. X-ray energy dispersive spectra were obtained in the same equipment operating at 15 KeV, and chemical elements were identified by their characteristic peaks.

#### Crystallinity standard obtained through X-ray diffraction

NPC and hydroxyapatite samples were deposited on silicon substrates, air dried, and analyzed in a polycrystalline diffractometer (Geigerflex-RIGAKU), comprising a copper pump. For each sample, accumulations were performed at 1-minute intervals at a reading angle 2θ with a step of 0.1°. The measurements were performed in the Crystallography Laboratory, Department of Physics, University of Minas Gerais. The acquired data were plotted using OrigimPro 8 SR0 program.

### NPCFF and doxorubicin association

#### Elaboration of standard curve for doxorubicin

Aqueous solutions of doxorubicin (100, 10 and 1 μg ml^-1^) and the stock solution (1mg/ml) were read in a final volume of 200μL in VARIOSKAN (Multi Reader Device - Thermo) in a wavelength range from 200 to 1000 nm. The absorbance values obtained for the solutions at the wavelength of 480nm were used to elaborate the standard curve (absorbance x concentration) for doxorubicin using the Origin Pro 8® software.

#### Preparation of doxorubicin-associated nanostructured phosphate compounds (NPCF-DOX)

The preparation of NPCF-DOX was obtained from aliquots of 20 μL of doxorubicin (1 mg ml^-1^). Each aliquot received increasing concentrations of NPCF (0.1, 0.2, 0.3 and 0.4 mg ml^-1^), and was eluted to the final volume of 200 μL using milli-Q water. After centrifugation at 5000rpm for 1 minute, the supernatants were separated and the resulting precipitates eluted in 200 μl of milli-Q water.

#### Calculation of the percentage of association between NPCF and doxorubicin

The association between doxorubicin (0.1 mg ml^-1^) and different concentrations of NPCF (0.1, 0.2, 0.3 and 0.4 mg ml^-1^) was calculated using the standard curve obtained for aqueous solutions of the drug. The regression equation and the linearity (r^2^) obtained were *y = 0*.*0039 + 0*.*0429x* and 0.9999 respectively. Since *y = absorbance and x = concentration of doxorubicin*, it was possible to calculate the concentration of drug present in washed NPCF-DOX samples and their respective supernatants. The calculation of the loading of the drug to the nanostructured matrix was made according to the equation:

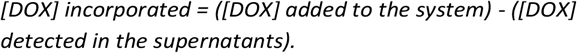

The percent binding of doxorubicin to NPCF was calculated based on the equation:

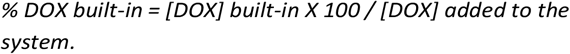

### Biological Assays

#### Cell Cultures

Human mammary tumor cells (MDA-MB-231 and MCF-7) were maintained in a humidified atmosphere with 5% CO2 at 37°C and culture using a Dulbecco’s Modified Eagle’s Medium (DMEM) medium (Sigma–Aldrich) supplemented with fetal bovine serum (FBS) (10%) (Cripion Biotechnology LTD), sodium bicarbonate (5 mM) (Chemical Kinetics LTD), penicillin (100 U ml^-1^), streptomycin (0.1 mg ml^-1^), amphotericin B (0.25 mg ml^-1^) (Sigma–Aldrich) and gentamycin (60 mg ml^-1^) (Schering–Plough). Prior to tests, exchanges of culture medium were performed at 48-hour intervals during the expansion period.

#### Detection of the CD44 expression in human mammary tumor cells MDA-MB-231 and MC7 via immunocytochemistry

Human mammary tumor cells—MDA-MB-231 and MCF-7— were plated into 48-well plates at densities of 2 × 10^4^ and 4 × 10^4^, respectively, based on the growth capacity of each lineage. After a 48-hour expansion period, cells were washed twice in PBS (0.15 M), fixed in paraformaldehyde (3.7%) for 20 minutes, washed in PBS twice, and permeabilized with Triton X100 (0.2%) for 10 minutes. After being washed with PBS three times, the cells were placed in a blocking solution (PBS + 1% BSA) for 30 minutes and incubated with polyclonal rabbit anti-CD44 ab41478 primary antibody (1: 250) in a block solution for 24 hours. Next, an incubation with Alexa555-labeled anti-rabbit secondary antibody (1:500) in a blocking solution lasted 1 hour, and at the end of this period, the cells were washed three times with PBS for 5 minutes. Nuclear labeling was done with Hoescht (1 μg ml^-1^, diluted in a blocking solution) for 1 hour at room temperature. At the end of the labeling procedure, the preparations were observed in OLYMPUS IX70 inverted fluorescence microscope using a 20× objective.

#### MTT Assay

The viability of the untreated MDA-MB-231 and MCF-7 strains exposed to NPCs was evaluated in 24 hours by assays using 3- (4,5-dimethylthiazol-2-yl) -2, 5-diphenyl tetrazolium bromide (MTT) (Cat. M-6494. Invitrogen). The assay is based on the reduction of the tetrazolium salt, in formazan crystals, by the dehydrogenase enzyme, which is active only in viable cell mitochondria.

For this purpose 10^5^ cells were seeded in 48-well plates, with a final volume of 400 μL of the DMEM medium (supplemented with 10% fetal bovine serum) per well. Cells were treated, in triplicate, with NPC, NPCF, and NPCFF at 1.0, 5.0, and 25.0 μg ml^-1^ concentrations. Samples used as positive control received no treatment, while other samples were treated with free rhodamin 6G at concentrations of 1.0, 5.0 and 25.0 μM. After each treatment, the culture medium was removed and 130 μL of fresh medium and 100 μL of MTT solution were added to each well. The samples were kept in a CO2 greenhouse for 2 hours. At the end of this interval, 130 μL of SDS (10% HCl) was added, and after 18 hours of incubation, the absorbance values of the samples were obtained in a spectrophotometer reader using a wavelength of 595 nm. ^35,36^

#### A CD44+ cell line (MDA-MB-231) real-time exposure to NPCFF

MDA-MB-231 cells maintained in culture under the conditions described above were plated on a 25 mm diameter sterile plastic dish with glass bottom cover. The cells had a density of 10^5^ cells in 1 mL DMEM (10% fetal bovine serum). After the 24-hour incubation period, the cells were treated with NPCFF (25 μg ml^-1^) and free rhodamin 6G (50 μM) Control samples were not treated.

Real-time image acquisition was made by using BioStation IM (NIKON), equipped with a video camera. The cells were observed for 30 minutes from the time of treatment addition, thereby capturing 30 images (1 photo per minute), with 4 fields each. The images were analyzed in the Image Pro Plus 4.0 ® program and the graphs (arbitrary fluorescence units as a function of time) plotted on GraphPad Prism®.

#### NPCF Internalization in Human Breast Tumor Cells (MCF-7)

MCF-7 cells were plated at a density of 2.5 × 10^4^ cells in a 1 mL (DMEM-10 % fetal bovine serum) in a sterile glass cover slip deposited in the bottom wells of a 48-well plate. After 24 hours of incubation, the cells were treated with NPCFF (60 μg ml-^1^) during periods of 10 and 20 minutes. At the end of the interaction, the culture medium was removed, and the cells were fixed in paraformaldehyde (4%) for five minutes. After removal of the fixator, cells were washed in PBS (10%) and permeabilized with Triton (X-100) for 5 minutes, and incubated with 488 phalloidin (30 μL) (Sigma–Aldrich) (PBS-BSA 1:40) for 1 hour. After further washing, they were incubated with Blue Dapi (30 μL) (Sigma–Aldrich) (PBS-BSA 1: 1000) for 1 minute. The entire procedure was performed in the dark. The cover slips were attached on the glass slide with a mounting medium (DDP) and observed through a confocal microscope with a digital camera that engages the microcomputer to obtain images with magnifications of 40× and 60×. The scanned images were processed in the program Image J.

#### Verification of the NPCFF internalization in human mammary tumor cells (MDA-MB 231 and MCF-7) through analysis of ultrastructure by scanning electron microscopy and elemental composition by x-ray microanalysis

For the determination of possible NPCFF internalization, MCF-7 and MDA-MB-232 cells were cultured on glass coverslips in the density of 1 × 10^5^ cells.ml^-1^ in DMEM supplemented with 10% FBS. After a 24 hour period for adherence and expansion in a greenhouse with controlled atmosphere (5% CO_2_) and temperature (37 °C), the cells were challenged with NPCFF (25 μg ml^-1^) during 24 hours. After the treatment, the samples were washed 3 times in PBS to remove the culture medium. The glass coverslips were then air dried, carbon double-tape mounted on aluminum stubs and vacuum metallized at 10^−5^ mBar, with Hitachi evaporator carbon (HUS4G). The processed samples were analyzed in the FEG Scan Electronic Scanning Electron Microscope (FIB - Quanta FEG 3D FEI), consisting of: a dual system with ionic and electronic beam, and a field emission electron gun. The resolutions of the apparatus were of 0.8 nm for the electronic beam and the focal length was of 3 to 99 mm in high and low vacuum. For the acquisition of images and analysis of the samples were used the secondary electron detectors, backscattered electrons and detector to obtain spectra of dispersive x-ray energy.

The images acquired by secondary electrons were used for the morphological description of the cell surface and confirmation of the structural integrity of NPCFF. With the images obtained by backscattered electrons were determined the sites of greater electro-density potentially containing NPCFF. For these analyzes we use electronic acceleration voltage between 5 and 15 kV. The electrodensity points selected in each sample were analyzed using a point electron beam with a minimum accumulation time of 90 seconds using the electronic acceleration voltage of 30 kV. The analysis described generated X-ray spectra whose peaks corresponding to chemical elements were identified, making possible the determination of the elemental composition of each analyzed point.

#### Evaluation of viability of human mammary tumor cells exposed to NPCF-DOX and free DOX

Cells of the MCF7 and MDA-MB 231 strains were seeded in 48 well plates at a density of 2 × 10^4^ and 1 × 10^4^ cells mL^-1^, respectively, in a final volume of 400μL DMEM medium (supplemented with 10% FBS) per well. Treatment with NPCF-DOX was performed using a stock solution prepared with 0.4 mg mL^-1^ NPCF and 0.1 mg mL^-1^ doxorubicin which was aliquoted so as to provide the drug at treatment concentrations of 0.01; 0.1; 1 and 10 μM. Treatments using free doxorubicin at concentrations of 0.001; 0.01; 0.1 and 1 μM and the positive control (without any treatment) were also performed. All treatments were performed in triplicate. After 24 hours of exposure, cell viability was assessed as described above.

#### Assessment of membrane disruption of MDA-MB231 and MCF-7 cells after NPCFF-DOX exposure

MDA-MB 231 and MCF-7 cells were seed (1 × 10^6^) in a bottle containing 5 mL of DMEM medium supplemented with 10% FBS. After 24 hours of exposure to NPCF-DOX (10 µM for both) and doxorubicin (0.1 µM for MDA-MB-231 and 1 µM for MCF-7) the cells that recevied no treatment and those with compromised membrane integrity were detected by a flow cytometric assay using propidium iodide staining. MDA-MB 231 and MCF-7 cells, in 100 μl aliquots were incubated for 15 minutes with 3μl of propidium iodide (50 μg ml^-1^) (PI—Sigma) in the dark. The samples were resuspended in 50 μl of PBS and immediately analyzed (without fixation) by flow cytometry using a BD FACScan flow cytometer (Becton Dickinson) using an air-cooled 488 nm argon laser. The maximum fluorescence excitation and emission values are 535 nm for PI. The propidium iodide fluorescence of cells after tratmentes was examined in 30,000 cells per sample. All FACS parameters (FSC and SSC) and region settings were kept identical throughout all experiments. The data were analyzed using FlowJo software®.

### Statistical Methods

Statistical analyses, when applied, were performed using GraphPad Prism 5.0 software (Prism Software, Irvine, CA, USA) using analysis of variance (two-way ANOVA) and followed by Bonferroni post-test. The results were then expressed as a mean standard error with 95% confidence interval. Results where p < 0.05 were considered statistically significant.

## Results

### Morphological and Physicochemical Characteristics of NPC and NPCF

NPC and NPCF were synthesized as described in experimental section. The zeta potential and conductivity of NPCs, NPCFs, polyethylene glycol (PEG), hyaluronic acid (HA) and the mixture of PEG and HA are shown in Table 1.

**Table 1:** Zeta potential and conductivity acquisition for NPCs and associated molecules.

| NPC/Molecule | Zeta Potential (mV) | Conductivity (mS cm<sup>-1</sup>) |
| --- | --- | --- |
| HA | -24.7 | 0.227 |
| PEG | -23.4 | 0.244 |
| HA+PEG | -27.9 | 0.395 |
| NPC | -23.0 | 0.0082 |
| NPCF | -14.2 | 0.1780 |

The nanoparticles zeta potential values remained negative after functionalization with the values of -23.0 mV (NPC) and -14.0 (NPCF). The conductivity increased from 0.0082 to 0.1780 mS cm^-1^. The functionalization process decreased the negative charge on the nanoparticles by approximately 38.2%.

The synthesized NPCs demonstrated surface roughened spherical shape, as depicted by scanning electron microscopy (SEM) images (Fig. 1A). The images of secondary electrons, acquired at 15 kV accelerating voltage, have allowed determining that NPCFs maintained their spherical shape and were found immersed in the organic fractions added to the system.

**Fig. 1:**
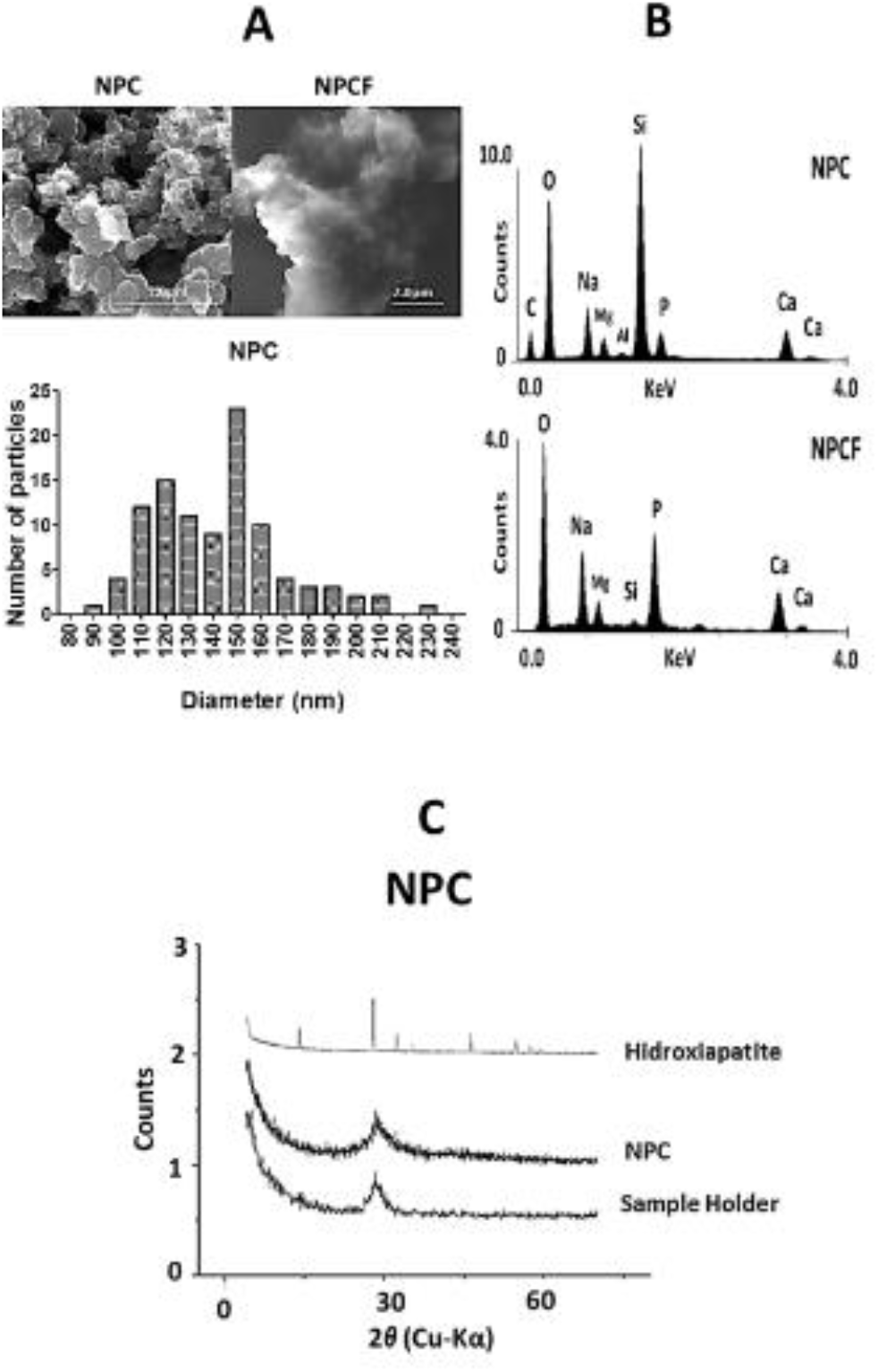
A. Scan electron microscopy photomicrographs of NPCs and NPC diameter distribution in nanometers. B. Energy dispersive X-ray spectroscopy (EDS) spectra obtained for NPC and NPCF showing their elemental composition. C. X-ray diffraction (XRD) of hydroxyapatite, NPC and the sample holder of the equipment. NPC and the sample holder present an amorphous pattern.

The mean diameter of NPCs was 149 nm; however, with a confidence interval of 95%, the diameter ranged between 136 and 147 nm (Fig. 1A).

### NPC and NPCF Elemental Composition through X-ray Microanalysis

The spectra obtained by analytical electron microscopy (Fig. 1B) depict characteristic peaks of C, Na, Al, and Si in NPC and NPCF. These are signals derived from the metallization process (C), stub (Al), and glass coverslip (Si) used as support for the samples. The other elements—Mg, Ca, and P—are characteristic of NPC inorganic matrix.

### NPC X-ray Diffraction (XRD) Patterns

The XRD patterns (Fig. 1C) provide the crystallographic natures of NPC, the synthetic hydroxyapatite (mainly because of crystalline calcium phosphate) and the sample holder device (amorphous). NPC has a single broad peak at 2θ = 28.3° corresponding to that obtained for the port silicon sample, thereby indicating its amorphous nature.

### Biological Assays

#### CD44 expression in MDA-MB-231 and MCF-7 Cell Lines

To confirm that the human mammary tumor cell lines used in this study express the gene for CD44, and were, therefore, potentially capable of interacting with nanoparticles functionalized with hyaluronic acid, immunohistochemistry for CD44 was performed on MCF-7 and MDA-MB-231 cells (as described in experimental section). Fluorescence microscope images demonstrated that both strains were positive for CD44 (Fig. 2A).

**Fig. 2:**
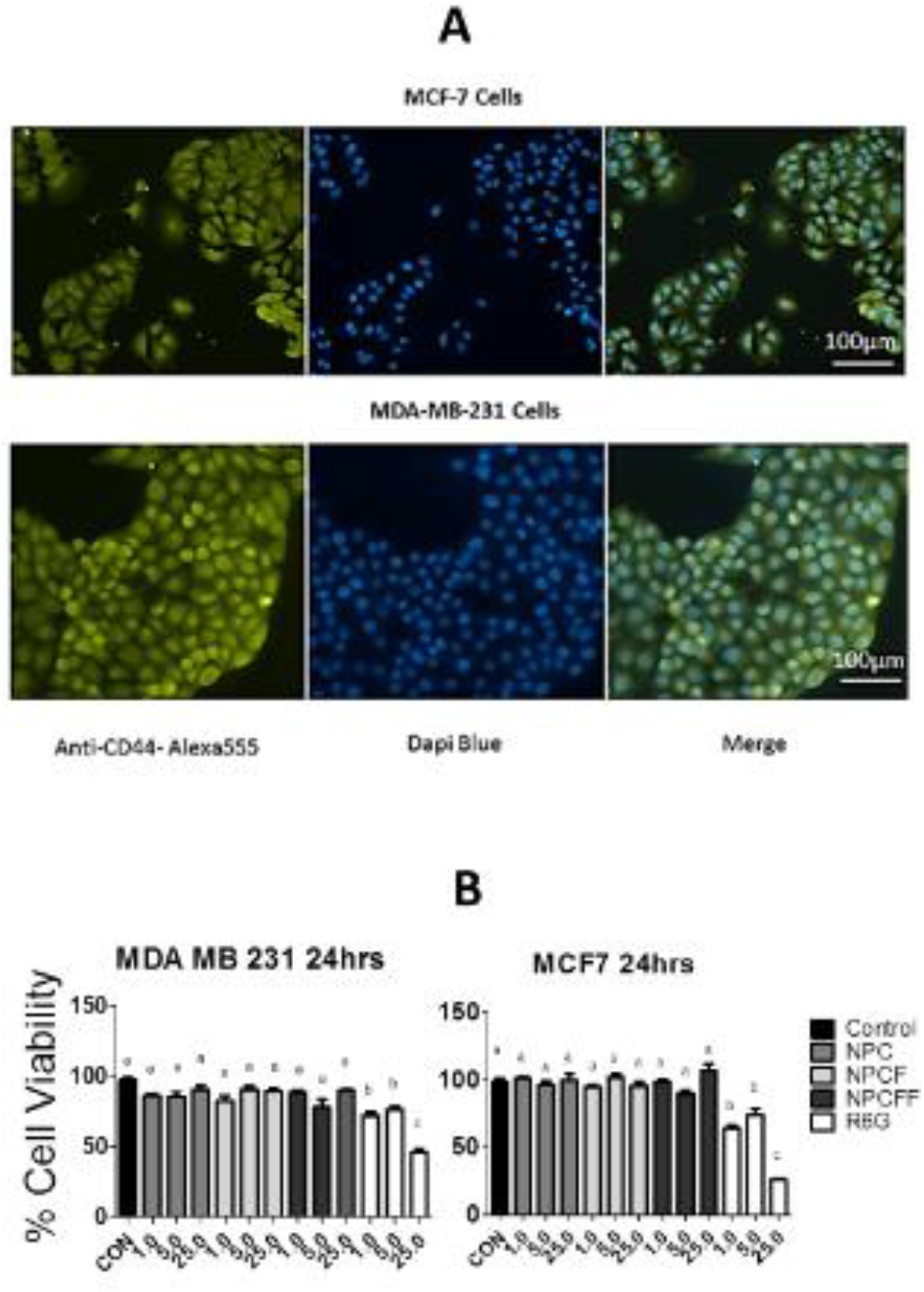
A. Immunocytochemistry for CD44 performed on MCF-7 and MDA-MB-231 cells. Fluorescence microscope images showed that both strains are positive for CD44 (green labeling). In blue, nuclear staining performed with Dapi Blue. B. Cell viability for MDA-MB-231 and MCF-7 cells treated with 1, 5 and 25 µg mL^-1^ NPC, NPCF, NPCFF and 1, 5 and 25 µM rhodamine 6G for 24 hours.

#### Toxicity of NPC, NPCF, NPCFF, and rhodamin 6G in MDA-MB-231 and MCF-7 Cell Lines

Treatment of human breast tumor cells— MCF-7 and MDA-MB-231—with NPC, NPCF, and NPCFF at increasing concentrations for 24 hours did not cause any cytotoxic effect when compared to no exposed cells (Fig. 2B).

Cells exposed to free rhodamin 6G at the same concentration for 24 hours demonstrated significant reduction of viability even in the lowest concentration. The cell viability reduction caused by the highest concentration of free rhodamin 6G was 75% in MCF-7 cells and 55% in MDA-MB-231.

These results show that the three types of phosphate-based nanoparticles, proposed in this study, do not show significant cytotoxic activity in either MCF-7 or MDA-MB-231. The functionalization with HA and PEG (NPCF) and the posterior connection with rhodamin 6G cells (NPCFF) did not induce cytotoxic effect. These findings are consistent with the fact that NPC and NPCF are not associated with any antimitotic compound, and are, by themselves, harmless to tumor cells in question. The association of rhodamin 6G, has proven to be not cytotoxic, wherein NPCFF appears to decrease the cytotoxic effect of the fluorochrome.

#### A CD44+ Cell Line (MDA-MB-231) Real-time Exposure to NPCFF

MDA-MB-231 cells exposed to NPCFF for 30 minutes and monitored in real time were also exposed to equivalent concentrations of free rhodamin 6G under the same conditions. The addition of free rhodamin 6G turns the cells into being medium fluorescent. Visually as well as quantitatively, the intensity of fluorescence in samples that received free rhodamin 6G was the same throughout the duration of the experiment (Fig. 3A). Cells exposed to NPCFF do not spontaneously become fluorescent, and the temporal progression of the fluorescence intensity in the cytoplasm can be visually noticed, thereby indicating their biologically mediated internalization, as suggested by electron microcopy analysis. Quantitatively, fluorescence increases markedly after 10 minutes of exposure, being superior to that detected in the rhodamine 6G treatment, after 15 minutes. Cells treated with NPCFF demonstrate no apparent signs of cell death at the end of the experiment, unlike in the case wherein cells were treated with rhodamin 6G, indicating morphological changes consistent with apoptosis signs.

**Fig. 3:**
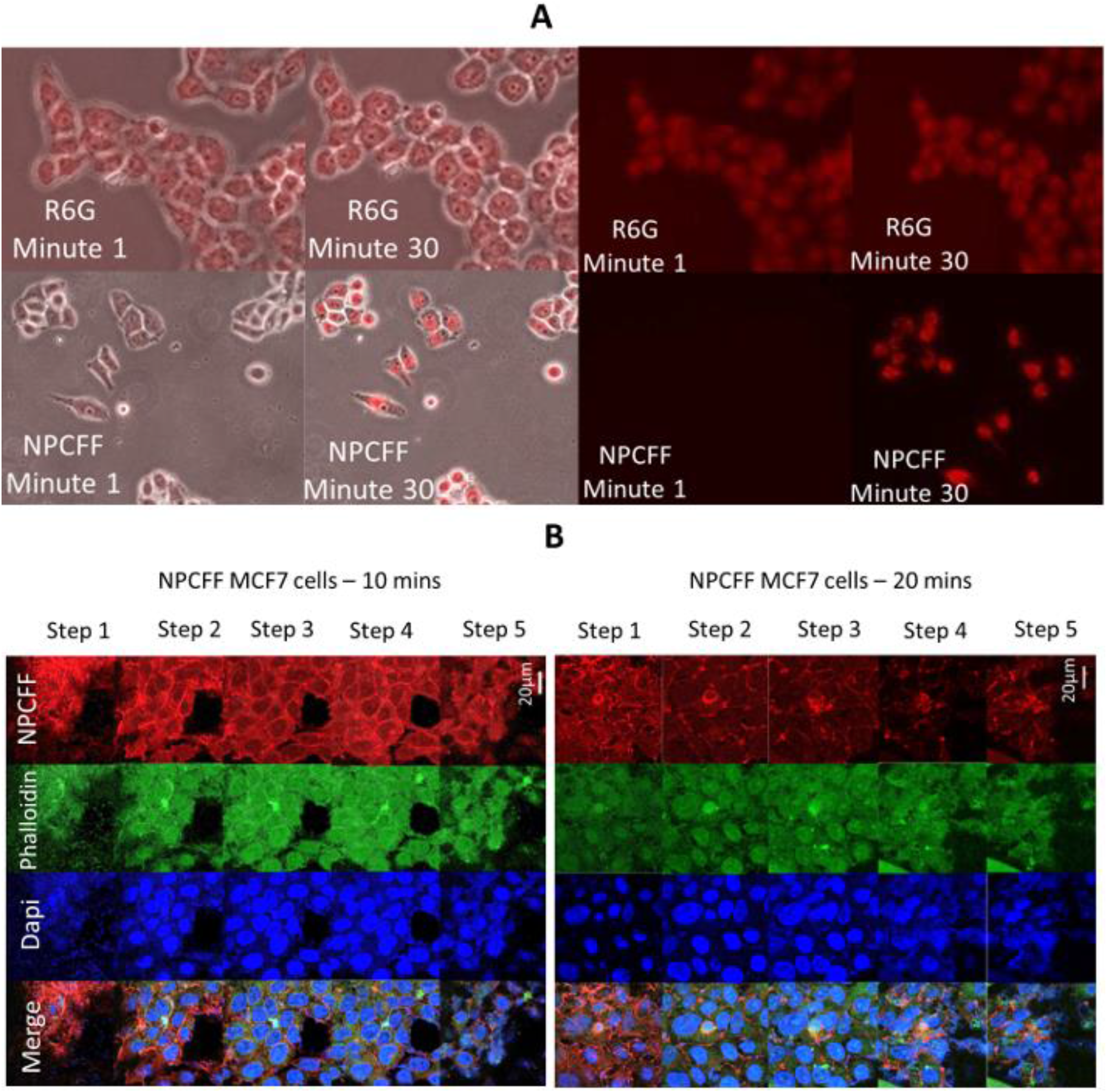
A. MDA-MB-231 cells exposed to free rhodamin 6G and NPCFF for 30 minutes and monitored in real time. Fluorescence quantification of MDA-MB-231 cells exposed to free rhodamin 6G and NPCFF for 30 minutes and monitored in real time. B. Human mammary tumour cells (MCF-7) staining with Blue DAPI (nucleus), 488 phalloidin (cytoplasm), and treated with NPCFF (R6G) for 10 and 20 minutes, respectively. Each step—0.25 µm in the Z-Series (Magnification 60× using confocal microscope).

#### NPCFF Internalization into a CD44+ Tumor Cell Line (MCF-7)

Conformal microscopy images of MCF-7 cells treated with NPCFF (60.0 μg ml^-1^) captured during exposures of 10 and 20 minutes demonstrate labeling of nuclear chromatin blue (Blue DAPI), cytoskeletal green (488 phalloidin), and NPCFF red (Fig. 3B).

The distribution of markings along the Z-axis allowed visualization of nuclear domains defined by the blue dye, green cytoplasm, and red NPCFF. No channel overlap was observed, demonstrating the absence of co-localization between the nanoparticles, cytoskeleton, and heterochromatin. At greater magnifications and for the same exposure times, it is possible to see the robust aggregation of nanoparticles (NPCFF) mainly in the peripheral region of the cell as well as in the cytoplasmic and nuclear domains. In the cytoplasm, they displayed a homogeneous distribution, despite the distance between the cell and the culture medium. Lower staining intensity was observed in the nuclear portion, manifesting mainly in the euchromatin regions. Our data suggest that NPCFF particles can be successfully adhered to and internalized in CD44 positive MCF-7 cells.

#### CD44+ Cell Lines (MDA-MB-231 and MCF-7) Fine Structure and Elemental Composition after Exposure to NPCFF

After interaction with NCPFF for 30 minutes, the samples containing MDA-MB-231 and MCF-7 cell surface demonstrated a rough and sunken aspect in the SEM images obtained by secondary electrons. NPCFF nanoparticles were present at various points in the field, adhered to the cell membrane surface with a spherical shape, and eventually being internalized. Through back-scattered electron images, the NPCFF electron density in a plane below the cell membrane confirmed this internalization. The NPCFF can be recognized as whitening spherical aggregates, which differ from crystal-shaped NaCl deposits (Fig. 4). Moreover the spectra obtained by X-ray microanalysis confirmed the elemental nature of the most electrondense structures such as the salt crystals, where Na and Cl ions are predominating and NPCFF, where an distinctive occurrence of P and Ca can be seen (Fig. 4).

**Fig. 4:**
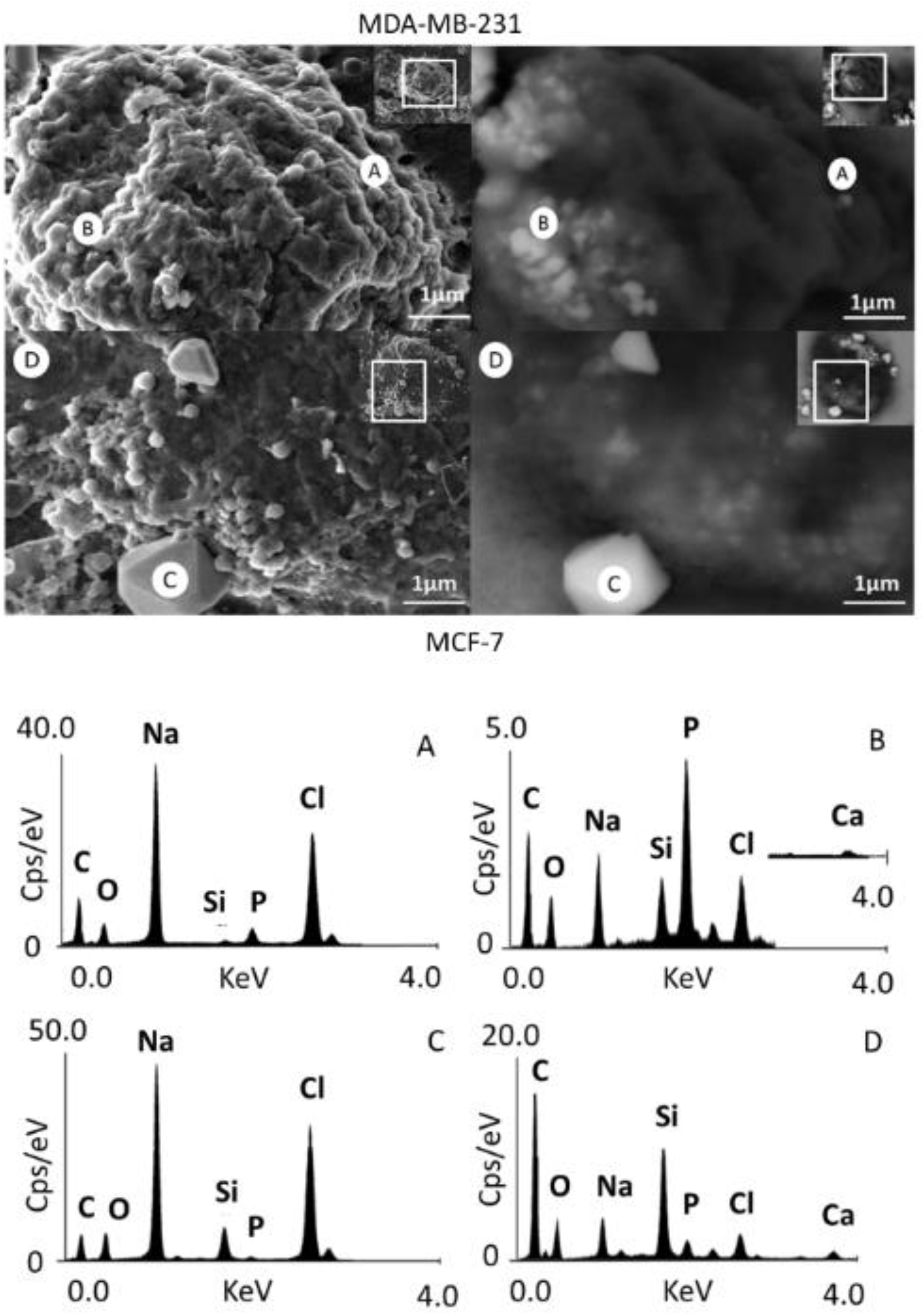
Secondary and back scattered scanning electron microscopy images of MDA-MB-231 and MCF-7 cells after 30 minutes of exposure to NPCFF (25 µg mL^-1^). A, B and C and D corresponds to the sites where the EDS were acquired.

#### Amount of doxorubicin binding to NPCF

The absorbance spectra obtained for the supernatants resulting from the NPCF-DOX wash were similar to the spectrum obtained for free doxorubicin (0.1mg ml^-1^), having high absorbance values with peak at 480 nm. The spectrum obtained for NPCF not associated to the drug presented a spectral profile similar to that of water, with no peak in the evaluated range. The values and peak of absorbance obtained for the NPCF-DOX samples decreased discretely with the increase of the NPCF concentration in the sample, being smaller in the sample containing NPCF at 0.4 mg mL^-1^ (Fig. 5A). This behaviour can be due to the high density of solid particles present in the most concentrated NPCF systems as a non-specific absorption form, more evident in the pure NPCF spectrum, since the concentration of DOX in the other samples is constant. The samples containing the washed precipitates recorded absorbance values markedly lower than those obtained for their supernatants and for free doxorubicin but maintained an evident absorbance peak at 480 nm. This indicates that after washing and centrifugation an aliquot of doxorubicin remained associated with NPCF, indicating the binding between the nanoparticles and the drug (Fig. 5A). The loads of the drug obtained in the NPCF-DOX samples containing increasing concentrations of NPCF are shown in Table 2. The most efficient association was in the NPCF 0.4 mg mL^-1^ sample which

**Fig. 5:**
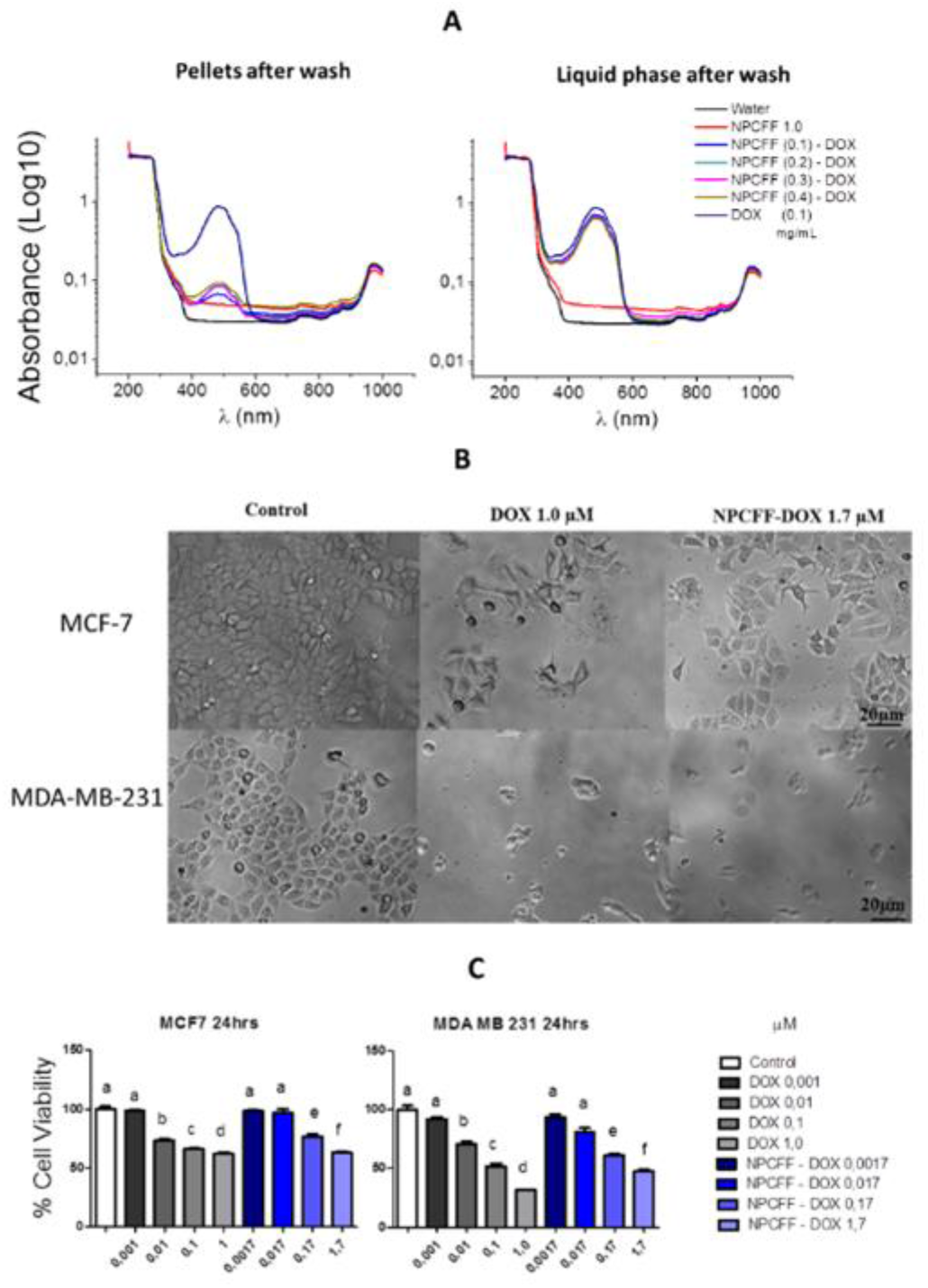
A. Absorbance spectra obtained for: supernatants obtained after NPCFF-DOX washing at the four nanoparticle concentrations (0.1, 02, 03 and 0.4 mg.mL^-1^), doxorubicin (0.1 mg mL^-1^), NPCFF (1.0 mg mL^-1^) and milli-Q water. Wavelength ranging from 200 to 1000 nm. B. Morphology of human mammary tumour cells MCF-7 and MDA-MB-231 after 24 hours of exposure to doxorubicin and NPCFF-DOX at concentrations of 1.0 and 1.7 µM. Inverted light microscope. Magnification: 10X. C. Viability of human mammary tumour cells treated with NPCFF-DOX and free doxorubicin.

**Table 2:** Load of doxorubicin in NPCF-DOX according to the nanoparticle concentration added to the system.

| [NPCF] in NPC-DOX mg mL<sup>-1</sup> | Load (%) |
| --- | --- |
| 0.1 | 5.95 |
| 0.2 | 10.15 |
| 0.3 | 13.08 |
| 0.4 | 17.72 |

#### Toxicity of NPCF-DOX in human mammary tumor cells MDA-MB-231 and MCF-7

The morphology of both cell types upon exposure to doxorubicin and CNF-DOX treatments is shown in Fig. 5B. The preconfluent appearance of untreated cells can be noted, whereas treatment with NPCF-DOX and free doxorubicin causes visible morphological changes in MCF-7 cells. The reduction of cell adhesion is observed, which makes its appearance more rounded with the apparent decrease in cell density. These changes are more evident in MDA-MB-231 cells when compared to MCF-7. MCF-7 cells underwent significant reduction in viability from the 1.0 μM drug concentration in NPCF-DOX. The decrease was dose-dependent, with a reduction in cell viability of 43% and 48% for treatments with doxorubicin at 1 μM and NPCF-DOX at 1.7 μM respectively (Figure 6). MDA-MB 231 cells treated strain NPCF-DOX containing 0.001 μM of doxorubicin did not present significant reduction of viability in relation to the control. The other treatments showed a significant reduction in cell viability, reaching 53% in cells treated with NPCF-DOX containing 1.7 μM doxorubicin (Fig. 5C).

**Fig. 6:**
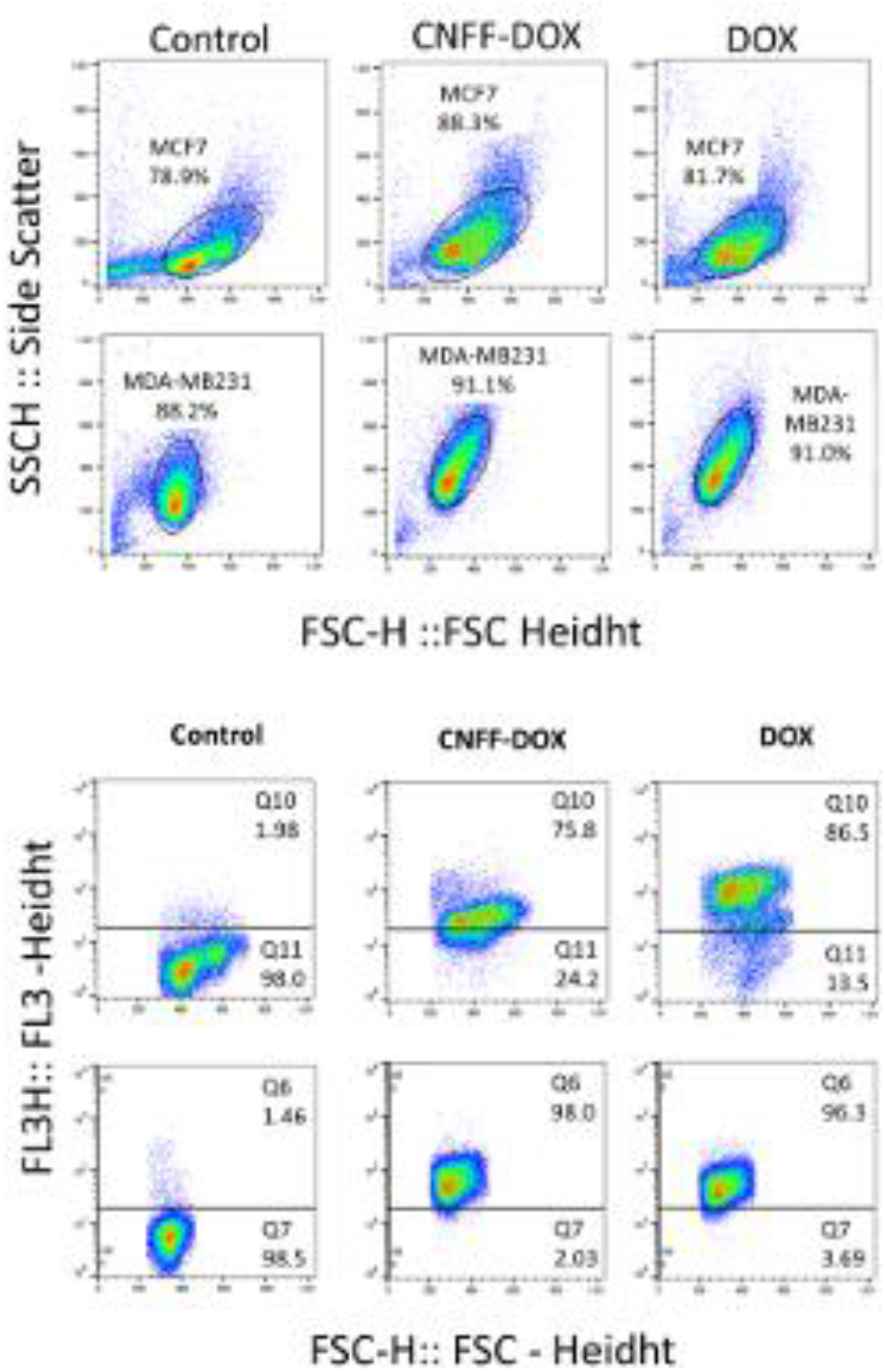
MCF-7 and MDA-MB-231 cells populations analised by ssc and fsc parametres showing the gate used for membrane integrity analisys in control, exposed to DOX and NPC-DOX treatments. MCF-7 and MDA-MB-231 cells populations analised by Pi fluorencence and FSC showing the % populations with loss of membrane integrity analisys in control, exposed to DOX and NPC-DOX treatments.

#### Assessment of membrane disruption of human breast cancer cells exposed to NPCFF-DOX and free DOX

The membrane disruption was detected in MDA-MB-231 and MCF-7 cells by PI stain and analyzed by flow cytometry. The SSC-FSC cels profiles are presented in Fig. 6. In both cells types (MCF-7 and MDA-MB-231 a shift on FSC and SSC profiles were noted. The cells exposed to DOX and CNF-DOX increase sligtly the SSC ligth. The positive Pi labelled cells were obseved in in all cells exposed to NPC-DOX and DOX reaching to 98,3% and 87,0% to MCF-7 and 98,3% and 96% to MDA-MB-231 respectively, indicating their prevalence on cellular membrane permeability commitment (Fig. 6).

## Discussion

Usually, a nanoparticle less than 100 nm in size is the most desirable for biological applications, especially for in vivo delivery and targeting. ^37^ A multi-functional nanoparticle, containing a pH-sensitive calcium phosphate-based shell and an amphiphilic gelatin matrix, demonstrated controllable and sequential release of doxorubicin and corcumin. These nanoparticles had an average size of approximately 150 nm, however, with increasing concentrations of the targeting agent (Amifostin) the eventual size of the nanoparticle was no less than 170 to 200 nm. ^38^ An independent endeavor using pegylated-triple-shell calcium phosphate nanoparticles^39^ demonstrated that single and double shelled nanoparticles had an average diameter of about 73 and 109 nm, respectively. The functionalized triple shell nanoparticles, containing doxorubicin along with a fluorescent dye (Dy550) had larger diameters (between 127 and 156 nm). NPC, used in this study had an average diameter (149 nm) consistent with that found in other efficient nanocarriers and/or probes.^40^

It is known that the functionalization process is responsible for promoting changes in the surface charge on nanoparticles. Citrate surface functionalization and pegylation are amongst the alternative procedures used for calcium phosphate nanoparticles, often increasing their electronegativity. ^40^ In NPC, functionalization leads to a decrease in its negative charge possibly due to HA interactions.

PEG surface modification has been widely used to improve the efficiency of tunable nanoparticles, especially, by increasing their lifetime circulation and preventing phagocytosis by reticular system cells. ^6^ In liposomes, PEG addition may be achieved through pre and post conjugation methods. ^41,42^ In NPC, a post conjugation method was performed, where PEG 400 was added to an aqueous NPC solution without ligand induction or other controlled conditions. The NPC–PEG biding was expected to occur through self-assembly, wherein the PEG chain length and surface density directly affect the PEG layer. ^43^ The volume of PEG 400 (3 µL) added to the NPC solution during the functionalization process was decided based on the fact that there exists a maximum molar percentage of PEG that can be incorporated into the system. ^6^ The addition of the PEG layer was also expected to increase the NPC colloidal stability; however, it was observed that in terms of the role PEG played in the kinetics of nanoparticles, minimum work had been done in this regard. ^6^

Additionally, it was expected that the use of a targeting ligand could increase the efficacy of calcium phosphate-based nanoparticles as CD44+ tumours targeting systems. Many kinds of nanoparticles have been conjugated to targeting moieties while maintaining colloidal stability. ^44^ A number of previous studies, focusing on the role of HA in target CD44+ cells, have demonstrated good results in reducing cell viability or tumour volume in many types of cancer. ^45^ Given the multiple biological functions of CD44 and HA, they are also linked to stem-cell associated proprieties. The CD44 receptor is a cancer stem cell (CSC) marker for a large number of tumour entities, including breast cancer. ^46^ Several studies have shed new light on mechanisms through which CD44 modulates the functional proprieties of CSC. This way, CD44 is considered an attractive targeting for a particular tumour cell population. ^47^

The broad use of HA to improve CD44+ cell targeting is explained by the fact that HA selectively binds to CD44. HA nanoparticles are concentrated on the cell surface and internalized. Additionally, the toxicity of these materials is often lower than free compounds like drugs or organic dyes. ^48^ An HA molecular mass of 500 kDa could bind up to 200 CD44 receptors; thus, upon bidding, HA– CD44 could be internalized. ^49^ Therefore, these data suggest that CD44 pathway could be considered as a good choice in cancer therapy, and NPC functionalization with HA could leads to its internalization in MDA-MB-231 and MCF-7 cells.

It is very important to emphasize that the HA–CD44 molecular system provides nanoparticles with specific targeting capabilities,^50^ thereby constituting a potential tool for cancer management.

Some of the most promising systems for tailored markers are hybrid nanoparticles, which are fabricated by embedding organic dye molecules in a solid particle, wherein the use of a non-cytotoxic calcium phosphate matrix provides an alternative delivery system, along with other existing inorganic or metallic nanoparticles. It is known that the encapsulation of organic fluorescent dyes could increase their fluorescence, protecting them from photo degradation and increasing the accuracy of analyses. ^34^ Nanoparticles encapsulating rhodamin WT, fluoresceins, and Cy3 amidite were used in intracellular imaging to detect melanoma and smooth muscle cells, with simultaneous drug delivery. ^51,52^ Cy3 amidite calcium phosphate-based nanoparticles were also used to determine the quantum efficiency of encapsulated dye molecules compared to those of the free dye. A study demonstrated an increase in this property because dye molecules were shielded from solvent effects. ^53^ Thus, studies like those mentioned above have demonstrated the in vitro versatility of dye-doped calcium phosphate nanoparticles with little or no cytotoxicity and opened an avenue for studies regarding NPC and rhodamin 6G association for similar purposes.

Rhodamin 6G (C_28_H_3_0N_2_O_3_HCl) is a highly photo stable dye with a very high quantum yield (nearly 0.95) ^54^ however, some studies have demonstrated its association with nanostructured matrices, ^55^ tending to reduce its degradation improving efficiency. Many issues require consideration while using rhodamin 6G in associated with nanomaterials, especially solids matrices, since aggregation between the dye and material may produce dimers that modify the absorption spectra and decrease the fluorescence quantum yield and decay time. ^56^ In water solutions, as used in NPCFF preparation, at a range of concentrations (up to 10^−5^ M), dimers are responsible for fluorescence quenching, and when a specific concentration (10^−4^ M) is attained, the solution does not demonstrate lasing capability anymore. ^57,58^ Against these findings, we did not observe rhodamin 6G characteristic alterations after association with NPCF. NPCFF showed absorption spectra similar to that found in free rhodamin 6G, and even after four successive washing processes, the signals found in NPCFF were detectable in subsequent biological assays. Using methanol or ethanol as the solvent medium greatly inhibits rhodamie 6G’s tendency to form dimers; however, the NPCFF preparation must be done in water, because alcoholic solutions alter its colloidal stability state, especially in the presence of HA.

Calcium and phosphate are both ions present in the circulating blood stream and cells in millimolar concentrations ^59^ but when this concentration increases rapidly and reaches values above those physiologically acceptable, it could lead to apoptosis. ^60^ As a result, the cytotoxicity of calcium phosphate-based nanoparticles has been investigated in in vitro assays in a number of different cell lines. ^61^ In human monocytes-derived macrophages (HMMs), hydroxyapatite nanoparticles demonstrated toxicity in concentrations above 250 µg mL^-1^, which suggested that interference with calcium homeostasis would be the primary cause of toxicity. ^62^ In contrast, other studies demonstrated that fluorescein doped calcium phosphate-based nanoparticles did not produce any cytotoxicity or change in metabolism in mouse stellate ganglia neurons as measured by calcium pumping currents. NPC and NPCF did not cause cytotoxic effects in CD44+ human breast cancer cells MDA-MB-231 and MCF-7. It can explained by the absence of the crystalline phases on its composition, and also, because NPCF does not contain any cytotoxic agent or antimitotic drug in its composition. On the other hand, NPCFF contains rhodamin 6G, which is a cytotoxic agent, and hence, it could be expected to promote cell death. However, NPCFF was demonstrated to be non-cytotoxic, like the other NPC formulations. Lastly, free rhodamin 6G caused cell viability reduction even in low concentrations (1 µM). The cytotoxic effect was amplified with an increase in rhodamin 6G concentrations added to the medium (5 µM and 25 µM). It is desirable that nanostructured systems designed for cancer imaging are non- or minimally toxic to target and normal cells. ^34^ The major organic dyes in free form, including rhodamin 6G are toxic to cells, what was confirmed by our experimental results. The association with NPC do not alters major optical proprieties of rhodamin 6G and is proven that NPC–rhodamin 6G binding caused reduced cell cytotoxicity. This way, NPCFF could be used as a less invasive and damage-free method of tumour diagnosis and targeting.

It is known that nanoparticles enter the cells by endocytosis ^63^ despite the fact that mechanisms underlying cross membrane translocation are yet not completely elucidated. Polyethyleneimine nanoparticles are internalized into the cytoplasm and they escape endosomal vesicles, being transported in a microtubule-dependent manner, into the perinuclear region within 10 minutes. ^64^ On the other hand, quantum dots remain associated with lysosomes and endosomal compartments for days. ^65^ Cationic polystyrene (PS) nanoparticles cause lysosomal rupture, mitochondrial damage, and oxidative stress, all of this within the first hour of nanomaterial addition. ^66^

HepG2 cells were incubated with carboxy-coated, yellow–orange polystyrene (COOH-PS [YO]) (50 nm), plain yellow–green polystyrene (plain-PS [YG]) (50 nm), and FITC-labeled silica nanoparticles (50 nm) for 1 hour, followed by confocal image acquisition. These nanomaterials strongly accumulated in the cytoplasm and nucleus in a diffused pattern. Fluorescence measurements through confocal nucleus sections confirmed that a substantial amount of these nanoparticles had accumulated in this region. This accumulation pattern occurred after one hour of cell– nanoparticle interactions, and it is possible to notice that cytoplasm accumulation occurs intensely than in the nucleus. ^67^

Additionally, a transcriptional activator protein (tat)-cross linked iron oxide nanoparticle (tat-CLIO) was attached to FITC and Cy3 (tat-(FITCH)-Cy3-CLIO) and rapidly internalized by HeLa cells, attaining 100% labeling within 45 minutes. The amount of label per cell increased linearly with time until 3 hours, and cells, which were loaded for 1 hour, retained 40-60% of FITC and Cy3 labels. The use of confocal microscopy demonstrated that both dyes had entered nuclear and perinuclear domains after tat-(FITC)-Cy3-CLIO labelling.^68^ After 3 hours of internalization on HeLa cells, fluorescent cationic and anionic calcium phosphate nanoparticles started to dissolve under acidic conditions in the lysosomes. Similarly, TATp-liposomes, presented a rapid and efficient translocation of cells migrating to the perinuclear zone. The 200 nm Rhodamine labeled TATp-liposomes, loaded with FITC-dextram, rapidly translocate to human breast adenocarcinoma cells (BT20). ^69^

NPCFF behavior could also be reckoned to correspond to these findings, since we have observed its rapid and efficient internalization on MCF-7 and MDA-MB-231 cells. NPCFF’s differential is the fact that we could find considerable labeling within few (10–20) minutes, according to real-time observation. It is, therefore, safe to assume that NPCFF is a part of the nanoparticles collection that reaches the nucleus in a shorter time span. The rhodamin 6G addition to the NPCF matrix resisted washing, fixing, and staining processes prior to confocal microcopy, which removed the excess, non-adhering fluorochrome from the nanoparticle surface.

Pioneering studies report that the association of these therapeutic agents with nanoparticles represents an effective alternative to overcoming the drawbacks of conventional delivery methods. This is because nanoparticles have a reduced size that allows their penetration through the tissue capillaries, with efficient accumulation of the drug at target sites in the body.^70^ Thus, the systemic toxicity of the drugs is reduced and the efficiency of the treatment is increased. ^71^ In paramagnetic iron oxide nanoparticles associated with doxorubicin, PLGA and an aptamer with affinity for murine colon cancer cells (C26), the mean loading of doxorubicin to the nanostructured matrix was 3% of the drug added to the system (concentration of 10mg.mL^-1^). ^72^ In contrast, in mesoporous silica nanoparticles functionalized with PEG and amino-diacetic acid, the loading of the same drug was 9.3% of the concentration used in the synthesis (0.1mg mL^-1^). ^73^ In NPCF-DOX, loading of doxorubicin was approximately 18% from an initial concentration of 0.1mg.ml^-1^. Compared to some recently described nanomaterials, NPCF-DOX had a higher association rate from a lower concentration of drug. Thus, the use of NPCF as a platform for the association of doxorubicin is a potentially efficient and economical method. Functional mesoporous silica nanoparticles with the CD44 monoclonal antibody (CD44 McAb) were associated with doxorubicin and tested on MCF-7 / MRD1 cells (considered multi-resistant). Concentrations above 1.83 μg mL^-1^ caused a decrease in cell viability of approximately 20%, and the concentration of 18.3 μg mL^-1^ led to approximately 50% cell death. These values, however, do not differ significantly from those obtained for free doxorubicin.^73^ NPCF-DOX had a greater cytotoxic effect on MDA-MB-231 cells when compared to the effect caused on MCF-7 cells. This goes according to the resistance reported for MCF-7 cells. As with most nanoparticles used for the same purpose, NPCF-DOX showed a dose dependent effect in both cell types tested, causing about 40% death in MDA-MB-231 cells at a concentration of 0.17 μM and 53% At 1.7 μM. In MCF-7, the same concentrations caused approximately 23 and 47% death. NPCF-DOX in this case is as efficient as other nanoparticles recently associated with doxorubicin for the treatment of tumor cells.

## Conclusion

Nanoparticle phosphate-based composite (NPC) possess a good diameter reproducibly, size, surface charge, and amorphous nature consistent with good and desirable cell CD44 positive surface interaction. NPC is a good platform for an easy HA and PEG functionalization, and this process’s product (NCF) can be attached to a red fluorescent organic dye, maintain its abortion characteristics and reducing its cytotoxic effect (NPCFF). Since NPCFF are rapidly (from 10 to 20 minutes) internalized into CD44+ human breast cancer cells, we can consider it as an efficient, non-toxic, low-cost, and less invasive nanomarker. NPCF has the ability to self-assembly with doxorubicin forming a NPCF-DOX compound that has compatible cytotoxicity pattern for its use in the targeting of CD44+ tumour cells, evidenced in this work with the MCF-7 and MDA-MB-231 lines. In the future, NPCFF and NCFF-DOX potential for in vivo experiments could be investigated.

## Acknowledgements

This study received founds by research the funding agencies Conselho Nacional de Desenvolvimento Científico e Tecnológico (CNPq 12/2016) and Pró-reitoria de Pesquisa da Universidade Federal de Minas Gerais (PRPq-UFMG 02/2017).

## Notes

### Competing Interest Statement

The authors have declared no competing interest.

